# Shrimp draft genome contains independent clusters of ancient and current endogenous viral elements (EVE) of the parvovirus IHHNV

**DOI:** 10.1101/2021.12.29.474436

**Authors:** Suparat Taengchaiyaphum, Prapatsorn Wongkhaluang, Timothy W. Flegel, Kallaya Sritunyalucksana

## Abstract

**Background:** Shrimp have the ability to accommodate viruses in long term, persistent infections without signs of disease. Endogenous viral elements (EVE) play a role in this process probably via production of negative-sense Piwi-interacting RNA (piRNA)-like fragments. These bind with Piwi proteins to dampen viral replication via the RNA interference (RNAi) pathway. We searched a draft genome of the giant tiger shrimp (*Penaeus monodon*)(GenBank record JABERT000000000) for the presence of EVE related to a shrimp parvovirus originally named infectious hypodermal and hematopoietic necrosis virus (IHHNV).

**Results:** The shrimp draft genome contained 3 piRNA-like gene clusters containing scrambled IHHNV EVE. Two clusters were located distant from one another in linkage group 35 (LG35). Both LG35 clusters contained multiple DNA fragments with high homology (99%) to GenBank records DQ228358 and EU675312 that were both called “ non-infectious IHHNV Type A” (IHHNV-A) when originally discovered. However, our results and those from a recent Australian *P. monodon* genome assembly indicate that the relevant GenBank records for IHHNV-A are sequence-assembly artifacts derived from scrambled and fragmental IHHNV-EVE. Although the EVE in the two LG35 clusters showed high homology only to IHHNV-A, the clusters were separate and distinct with respect to the arrangement (i.e., order and reading direction) and proportional content of the IHHNV-A GenBank records. We conjecture that these 2 clusters may constitute independent allele-like clusters on a pair of homologous chromosomes. The third EVE cluster was found in linkage group 7 (LG7). It contained EVE with high homology (99%) only to GenBank record AF218266 with the potential to protect shrimp against infectious IHHNV.

**Conclusions:** Our results suggested the possibility of viral-type specificity in EVE clusters. Specificity is important because whole EVE clusters for one viral type would be transmitted to offspring as collective hereditary units. This would be advantageous if one or more of the EVE within the cluster were protective against disease caused by the cognate virus. It would also facilitate gene editing for removal of non-protective EVE clusters or for transfer of protective EVE clusters to genetically improve existing shrimp breeding stocks that might lack them.

## BACKGROUND

Non-retroviral viral gene sequences inserted into host genomes during the course of evolution began to be called endogenous viral elements (EVE) upon their discovery in vertebrate genomes [1]. However, evidence for EVE in animals was first reported for insects in 1999 [2], but the significance of the paper was not fully appreciated at the time of its publication. For shrimp, an EVE was described much later [3] but it was named “non-infectious IHHNV”.

In 2009, it was hypothesized that EVE (at the time called non-retroviral viral genome inserts) resulted from a mechanism that provided for heritable immunity in shrimp and insects [4, 5]. In brief, the mechanism was proposed to involve host recognition of invading viral mRNA followed by its use as a substrate for host reverse transcriptase (RT) to produce variable fragments of viral copy DNA (vcDNA). The vcDNA fragments would subsequently be inserted into the shrimp genome where they would produce antisense RNA that could induce an RNA interference (RNAi) response. This would lead to inhibition of viral replication and allow the host to accommodate one or more viruses (i.e., allow viral accommodation) in persistent infections without signs of disease. Tests for predictions of the viral accommodation hypothesis proceeded much faster with insects than shrimp. By 2020, the basic predictions regarding EVE were proven for mosquitoes [6–8]. The studies in both *Drosophila* and mosquitoes revealed additional detailed mechanisms for an unpredicted, specific adaptive response involving the ability of vcDNA to produce siRNA resulting in an immediate, adaptive cellular and systemic antiviral RNAi response [7, 9, 10]. For details see a recent review [5].

One of the insect publications [10] revealed that the vcDNA produced by host RT in response to RNA virus infection came in two forms, one linear (lvcDNA) and one circular (cvcDNA). The latter could be specifically isolated relatively easily and was shown, when injected, to protect insect hosts againt the homologous virus. Following the protocols described for cvcDNA isolation in insects, it was shown [11] that IHHNV-cvcDNA could be extracted from IHHNV-infected *P. monodon* and that it could inhibit IHHNV replication in whiteleg shrimp *P. vannamei* challenged with IHHNV prepared from the infected *P. monodon*. During sequencing of the cvcDNA extract, it was also found that some of the cvcDNA constructs had high homology (98-99%) to the GenBank records DQ228358 and EU675312 known to arise from the host *P. monodon* genome (i.e., from EVE) originally called “non-infectious IHHNV Type-A” (IHHNV-A) [3, 12]. Thus, the 2021 report [11] revealed that EVE can also give rise to cvcDNA, and it was hypothesized that the cvcDNA arose via EVE-produced RNA interacting with host reverse transcriptase.

Around the time that the work with IHHNV cvcDNA [11] was being done, a draft whole genome sequence (WGS) of a Thai *P. monodon* specimen was published [13], and we became interested to determine whether the WGS might contain EVE related to IHHNV. We considered this possible because the specimes we used for our cvcDNA work and the specimen used for the WGS project originated from the same domesticated shrimp breeding stock.

## RESULTS AND DISCUSSION

### Clusters of IHHNV-EVE were found in linkage groups 35 and 7

A general BlastN search of the Thai *P. monodon* WGS project (GenBank accession no. JABERT000000000) [13] using the queries GenBank record DQ228358 for non-infectious IHHNV-A and GenBank record AF218266 for infectious IHHNV confirmed the presence of 3 clusters of EVE derived from IHHNV (**Fig. 1**). Two of these IHHNV clusters (**Fig. 1A**) were located in linkage group 35 (LG35) and showed high sequence homology (98-99%) to IHHNV in GenBank accession numbers DQ228358 and EU675312. The other cluster (**Fig. 1B**) was located in linkage group 7 (LG7) and showed high sequence homology (98-99%) to GenBank accession number AF218266 for an extant type of infectious IHHNV. All three EVE clusters were clearly demarked by bracketing, direct host repeat sequences marked by red arrows in Fig. 1.

**Figure 1.**
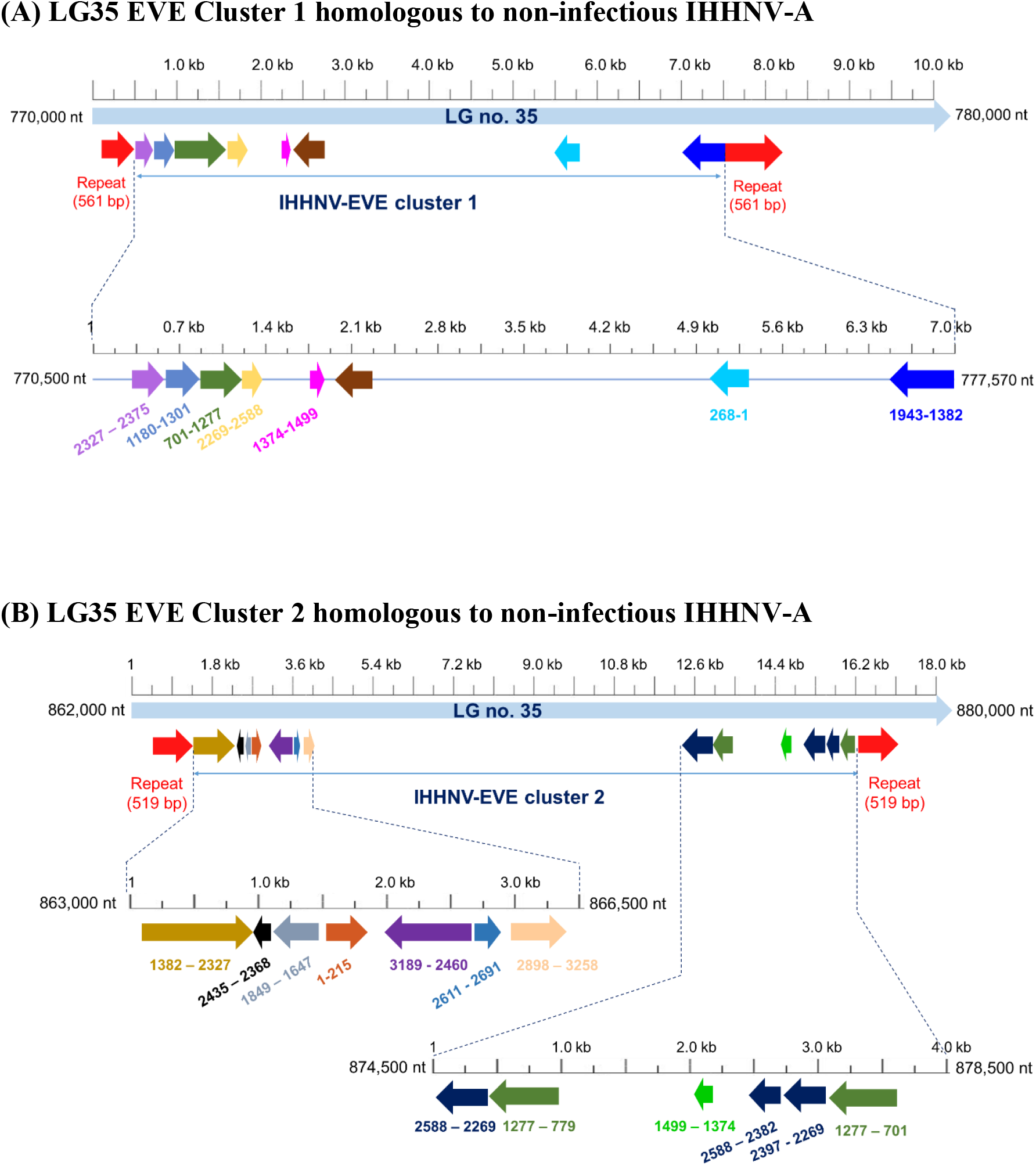
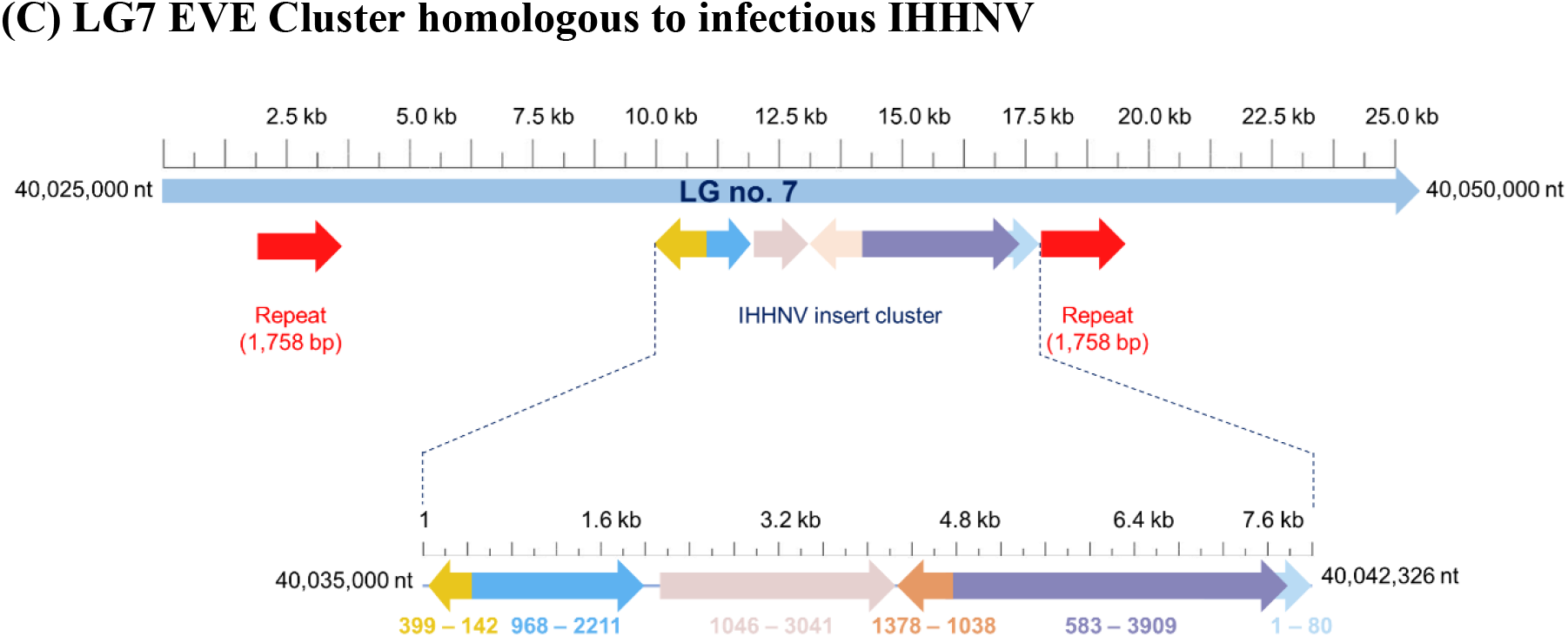
Schematic diagrams of IHHNV-EVE clusters in the draft *Penaeus monodon* WGS. **(A)** and **(B)** Sequence diagrams for Cluster 1 and 2, respectively, in LG35 with high homology to the non-infectious IHHNV-A query portion (1-3025 bp) of the GenBank record DQ228358. Some of the EVE sequences in the 2 clusters correspond to the same region of DQ228358 (i.e., the same color) but may differ in length and in reading direction indicated by arrowheads. Others are unique to each cluster. A zoom-in expansion of each EVE cluster is shown beneath. Colored numbers below indicate the nucleotide positions corresponding to GenBank record DQ228358. The portion of the record related to the host transposable-element portion of DQ228358 is indicated by a dark brown arrow. (**C)** Diagram of the IHHNV-EVE cluster in LG7 (GenBank accession no. JABERT010000007. 1) with EVE showing high sequence identity (99%) to GenBank record AF218266 for infectious IHHNV. The numbers below the arrows represent the matching positions AF218266.

### Characteristics of EVE homologous to non-infectious IHHNV-A in LG35

The GenBank records for DQ228358 (in *P. monodon* from East Africa) and EU675312 (in *P. monodon* from Australia) were initially referred to as “non-infectious IHHNV Type A [3, 12] even though the authors knew that the sequences originated from the host shrimp genomes (*P. monodon*). The IHHNV-related portions of the records for DQ228358 and EU675312 share 99% sequence identity, indicating that they arose from a similar, ancient type(s) of IHHNV that is distantly related to the types of IHHNV currently reported to cause disease in cultivated shrimp [14]. We now refer to the two GenBank records DQ228358 and EU675312 as EVE of non-infectious IHHNV-A. Here we use the DQ228358 sequence for or comparative analysis but focus primarily on the portion of the sequence (1 to 3025 bp) that has homology to IHHNV. The remainder of the record (3026-4655 bp) constitutes shrimp host repeat and transposable element sequences.

We were not surprised to find EVE of IHHNV-A in the Thai *P. monodon* WGS database. This was because we had already reported the occurrence of a variety of cvcDNA sequences with high sequence identity (98-99%) to non-infectious IHHNV-A in *P. monodon* specimens obtained from a different generation of the same breeding stock from which the Thai genome specimen was obtained [11]. Those cvcDNA sequences covered 68% (779 – 3930 nt) of the matching portion of the DQ228358 record (including some transposable element portion) and could be assembled into a single linear construct. However, at the time of that publication, we did not know the actual composition of the genome region(s) from which the cvcDNA sequences arose. All we could conclude at the time was that a variety of IHHNV-cvcDNA types were produced, some linked with host transposable element sequences and some not.

Our search of the Thai WGS revealed that LG35 did not contain a single continuous sequence with high sequence identity to IHHNV-A. Instead, it contained many high-identity EVE fragments of variable length that were scrambled with respect to proportion, position and reading direction in the non-infectious IHHNV-A reference sequences. These scrambled EVE were arranged in LG35 in two distinct clusters separated from one another by more than 80,000 bp (**Figs. 1A and 1B**). The individual EVE are shown in different colors to allow easy visual comparison of the EVE to matching regions of the IHHNV portion of the DQ228358 reference sequence. The positions of the EVE in LG35 are shown below each cluster diagram in black typeface while the positions relative to the DQ228358 record are shown above in colors to match those of each EVE. The colored vertical lines indicate the boundaries of the individual EVE. The number and lengths of the individual EVE and the gaps between them in the 2 clusters in LG35 are shown in **Table 1**. The EVE in the two LG35 clusters contain some common DQ228358 coverage, but they are distinct from one another in terms of the EVE types, lengths and reading directions when compared to the DQ228358 reference sequence (Figs. 1A and 1B). The lengths of the EVE clusters bracketed by their respective repeat sequences were 6,744 bp for Cluster 1 is and 14,993 bp for Cluster 2.

**Table 1.** Lengths of the EVE and intervening sequences in the 3 clusters found in the draft WGS of Thai *P. monodon*.

| LG35 Cluster 1 related to DQ228358 |  |  | LG35 Cluster 2 related to DQ228358 |  |  |
| --- | --- | --- | --- | --- | --- |
| EVE no. | Size (bp) | Gap (bp) | EVE no. | Size (bp) | Gap (bp) |
| 1 | 48 | 8 | 1 | 945 | 2 |
| 2 | 118 | 3 | 2 | 67 | 8 |
| 3 | 576 | 0 | 3 | 202 | 234 |
| 4 | 319 | 348 | 4 | 729 | 189 |
| 5 | 125 | 128 | 5 | 215 | 14 |
| 6 | 204 | 2923 | 6 | 80 | 194 |
| 7 | 267 |  | 7 | 360 | 8125 |
| Mean | 237 | 568 | 8 | 498 | 0 |
| SD | 176 | 1161 | 9 | 319 | 1169 |
|  |  |  | 10 | 125 | 521 |
| LG7 Cluster 3 related to AF218266 |  |  | 11 | 128 | 43 |
| EVE no. | Size (bp) | Gap (bp) | 12 | 206 | 15 |
| 1 | 257 | 0 | 13 | 576 |  |
| 2 | 1243 | 101 | Mean | 342 | 876 |
| 3 | 1995 | 2 | SD | 272 | 2308 |
| 4 | 340 | 0 |  |  |  |
| 5 | 3326 | 0 |  |  |  |
| 6 | 80 |  |  |  |  |
| Mean | 1207 | 21 |  |  |  |
| SD | 1268 | 45 |  |  |  |

The unexpanded diagrams for Clusters 1 and 2 in Fig. 1A and 1B shared some structural features. For example, both of the 2 EVE clusters (Fig. 1A and 1B) were bracketed by a pair of host, direct-repeat sequences (red arrows) of 2 x 561 bp in Cluster 1 Fig. 1A) and 2 x 519 bp in Cluster 2 (Fig. 1B). In Cluster 1, the two repeats shared 98% sequence identity (547/561 bp) (Supplementary Fig. S2) while those in Cluster 2 shared only 77% sequence identity (400/519 bp)(Supplementary Fig. S3). These bracketing direct repeat sequences (red arrows) demarked the boundaries of the non-infectious IHHNV-A EVE clusters. They matched many other regions in several linkage groups of the Thai WGS database but at relatively low sequence identities (around 80% or less).

Within the boundaries of Cluster 1 there was a host retrotransposon element (dark brown arrow) of 561 bp (Fig. 1A) that had 98% sequence identity to a matching portion of the 1395 bp host retrotransposon sequence that is part of GenBank record DQ228358. However, there was also a hit for 1354/ 1395 bases (98% identity) for the same retrotransposon portion of the DQ228358 sequence beginning very distantly from Cluster 1 at position 23,321,662 in LG35. All other spaces that separated the EVE fragments in Cluster 1 showed homology to various other shrimp host retrotransposon elements or host repeat sequences (not shown).

In contrast to Cluster 1, the pattern for Cluster 2 in LG35 (Fig. 2B) did not include any retrotransposon sequence with high homology to that in GenBank record DQ228358 within the bracketing 519 bp direct repeat sequences (red arrows). Like Cluster 1, all sequences separating the EVE showed homology to a variety of other shrimp host retrotransposon and repeat sequences (not shown). The bracketing sequences for EVE Clusters 1 and 2, matched many other regions in several linkage groups of the Thai WGS database but at lower sequence identities of around 80% or less (not shown).

It is not known whether these two clusters in LG35 are linked on one chromosome of a diploid pair or whether they are matching “allele-like clusters” (“cluster-alleles”) located one each in the chromosome pair represented by LG35. Now that we have the sequences of the two clusters in LG35, it will be possible to design specific primers to screen for each type in the current generation of the original *P. monodon* stock. By identifying one specimen positive for both clusters and mating it with one negative for both, obtaining about half the offspring each carrying only one or the other cluster would confirm whether or not they are cluster-alleles. Finally, the forms of these 2 clusters resemble those described from mosquitoes as piRNA-like gene clusters containing fragmented sequences from mosquito RNA viruses [6, 7].

A preprint [15] using DNA extracted from an Australian *P. monodon* specimen for genome analysis revealed one EVE cluster in their Scaffold group 97 (SG97) with high identity to GenBank EU675312 (i.e., the Australian version of IHHNV-A with 99% sequence identity to DQ228358). Given the information from our earlier publication and from the work reported herein and from Australia, it is likely that the continuous sequence records for DQ228358 and EU675312 at GenBank are assembly artifacts, and that they were obtained from fragmented and scrambled target sequences that had sufficient overlap to result in their assembly into single linear sequences when using the sequence of infectious IHHNV (GenBank record AF218266) as a reference [3, 12].

The preprint of Huerlimann et al. [15] gives detailed analyses of the EVE cluster found on SG97 of their Australian *P. monodon* specimen plus detailed comparison of it to the two EVE clusters that they (like us) discovered in LG35 of the Thai *P. monodon* WGS. Thus, those interested the detailed comparison should consult the Australian preprint.

### The IHHNV-EVE in LG7 showed high sequence homology to infectious IHHNV

Of greatest interest to us was the EVE cluster located in LG7 **(Fig. 1 C)** that showed high sequence identity (95-99%) only to GenBank accession number AF218266 from an extant form of infectious IHHNV [17]. In other words, this cluster contained no EVE with high sequence identity to GenBank records DQ228358 and EU675312 (i. e., “ non-infectious IHHNV Type A) that were found in EVE Clusters 1 and 2 in LG35. However, the general, overall pattern for EVE Cluster 3 was similar to those of the two for IHHNV-A EVE clusters located on LG35. For example, the LG7 EVE were scrambled in terms of fragment location, portion and reading direction with respect to the AF218266 reference genome. They were also bracketed by two host direct-repeat sequences (red arrows) of 1,758 bp (98% sequence identity). The EVE were either contiguous or separated by host retrotransposon or repeat sequences. Again, the overall arrangement resembled the Pi-RNA-like gene clusters reported for EVE of RNA viruses in mosquitoes [6, 7].

Our work with IHHNV-cvcDNA [11] and the work with mosquitoes [7, 8], suggest that negative sense RNA transcripts of the IHHNV-EVE in this cluster would have the potential capability of inducing a host RNAi response against infection by homologous types of infectious IHHNV. If our conjecture above that the 2 IHHNV-EVE in LG35 are alleles turns out to be correct, the fact that there is only one IHHNV-EVE cluster in LG7 suggests that it might be a single cluster/allele with no matching IHHNV-EVE on the homologous chromosome. If this is so, it would provide an opportunity to prove whether or not the IHHNV-EVE in LG7 is protective against infectious IHHNV.

For example, primers could be designed to screen for the IHHNV-EVE cluster in LG7 in the current generation of the *P. monodon* stock. Several cluster-positive individuals could be mated with several cluster-negative individuals. If the cluster behaves in an allele-like manner, subsequent screening of the offspring from these crosses would reveal that at least one of the positive parents carried a single copy of the LG7 cluster/ allele and would yield half the offspring positive for the allele and half negative for it. The offspring could then be challenged with infectious IHHNV followed by qPCR to determine infectious IHHNV loads in shrimp with and without the EVE cluster. A significantly lower mean IHHNV load in the cluster-positive offspring compared to the cluster-negative offspring would indicate that the cluster was protective. If so, the stock owner would then be able to use PCR to select crosses to maintain the protective cluster in subsequent stock generations.

### Detailed analysis of the IHHNV-EVE in LG7

Altogether, the IHHNV-EVE cluster in LG7 (**Fig. 1B**) is 7,267 bp in length and contains 6 EVE having high sequence identity with a current type of infectious IHHNV (GenBank record AF218266. The EVE range from 80 to 3326 bp in length (**Table 1**) and are scrambled with respect to portion, position and reading direction when compared to the IHHNV reference genome AF218266. In comparison to the two EVE clusters for non-infectious IHHNV in LG35, 3 of the EVE in Cluster 3 are much longer and the gaps between the EVE are shorter or do not exist. The significance of these differences is currently unknown. However, the general pattern of EVE scrambling, bracketing by host repeat sequences and separation or not by host transposable element or repeat sequences is similar that in the other 2 EVE clusters in LG35.

### Trade consequences from shrimp stocks carrying an LG7 protective IHHNV-EVE

We carried out a Blastn analysis of the IHHNV-EVE cluster in LG7 against the target sequences for the PCR detection method (“309”) recommended for specific detection of infectious IHHNV in the OIE diagnostic manual [18]. The 309 method was designed to be specific for detection of infectious IHHNV [12]. Our analysis of the EVE cluster in LG7 revealed that 3 of the EVE (2, 3 and 5) contained sequences with 95% identity and 100% coverage for the target of the 309 method based on the AF218266 reference sequence (**Table 2**). However, despite the less than 100% overall sequence identities of the potential EVE targets, the primer sequences for the 309 method matched 100% with the relevant target sequences in all 3 of the potential EVE targets (**Supplementary information 1**). Thus, the potential amplicons would be equal length (indistinguishable) from those that would arise from infectious IHHNV and for the specimen used for the genome project would constitute a false positive test results for infectious IHHNV.

**Table 2.**
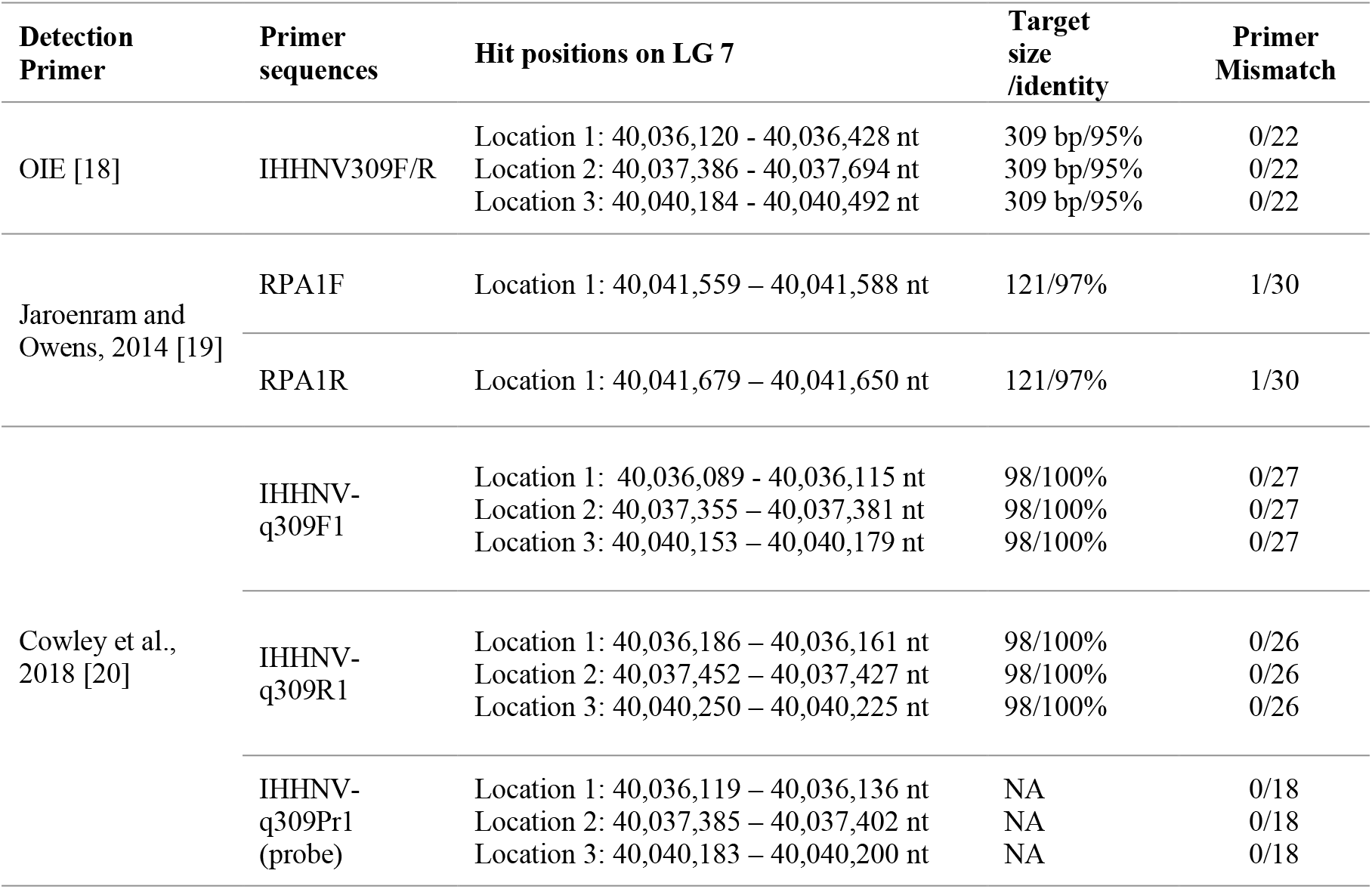
Potential targets in the IHHNV-EVE cluster of LG7. See the supplementary information for detailed information on the BlastN results.

In addition to the recommended method in the OIE manual, there are two published methods proposed to aid in the detection of infectious IHHNV while avoiding false positives arising from IHHNV-EVE. One of these [19] is an isothermal PCR detection method focused on a part of the infectious IHHNV genome that appeared to be of low prevalence or absent from IHHNV-EVE revealed by screening *P. monodon* from Australia. Our analysis of the target sequence for the primers from this method in the LG7 EVE for infections IHHNV (Table 2) revealed only one target sequence in the cluster in which there was a single base mismatch for each primer. There was a low calculated effect on the binding efficiency of the primers, so it is hard to predict whether or not the method would give a false positive test for infectious IHHNV with the potential LG7 target. In contrast, a more recent qPCR method designed to avoid false positive test results for infectious IHHNV that arise from IHHNV-EVE [20] would not be effective with the LG7 EVE cluster because there are 3 target sequences with 100% match to the primer and probe sequences.

In the past, without knowledge of EVE, shrimp specimens giving positive PCR test results with the methods above would have been considered IHHNV-infected and the positive shrimp specimens would have been discarded during the screening process to develop specific pathogen free (SPF) breeding stocks. In this way, it is possible that potentially protective EVE were discarded during stock development. This problem of false positive test results for infectious IHHNV arising from EVE has been raised previously [14, 21].

In addition, it is likely that shrimp products carrying the LG7 cluster would give false positive test results for the presence of infectious IHHNV, and this might result in the rejection of exported shrimp broodstock, shrimp PL and frozen shrimp products by countries that use the target of the OIE-recommended PCR method to screen imports for the presence of infectious IHHNV. It might be argued that use of a PCR method to detect such EVE followed by breeding selection to eliminate them from breeding stocks would be the simplest way to avoid this problem. However, such an approach might also remove naturally evolved, heritable resistance to IHHNV infection and lead to production problems for shrimp farmers.

In addition to the above potential for false positive results, another possibility has previously been raised [21]. This might occur in an SPF stock that had been developed by use of an internationally standardized PCR method to confirm absence of a particular virus based on a mutually agreed, small fragment of the target virus genome. This process would result not only in the discard of infected shrimp but also shrimp that carry EVE containing the PCR target sequence. Since some EVE may be protective and others not, it would be wise to choose the target region of any standard method to be in a genome sequence known to have no or low potential as a protective EVE.

However, this may not solve all the problems because individual shrimp in the stock population would carry a variety of difference EVE cluster/ alleles for a particular virus. Thus, it is possible that 2 individuals in a stock population might each carry one of 2 EVE in different cluster/ alleles but in the same reading direction and have the potential to re-establish the PCR detection target by crossover. Specifically, the separated EVE fragments in the two shrimp would have to have sequence overlap but with the EVE in one individual containing the forward 5’ primer target sequence for the chosen standard method but lacking the 3’ primer sequence, while the other individual would contain the 3’ primer sequence but lack the 5’ sequence. These EVE would escape the screening process to remove stock individuals with EVE that carry the target for the standard PCR detection method chosen. However, at some unpredictable time in many crosses and generations, these to sequences might end up as cluter/ alleles in a single individual by random assortment of chromosomes from its parents. If so, crossover events might occur in the overlap region to re-establish the PCR target sequence and give rise to a small portion of PCR positive individuals (i.e., “ pop-up” positive individuals) in the offspring of such individuals. This could happen even in a population of shrimp with a good history of freedom from the target virus [21]. For these reasons, it is essential that development of standard PCR detection methods involve a process of investigation and consultation among regulatory agencies and the companies or agencies that develop and maintain SPF shrimp breeding stocks for shrimp farmers.

### The evolutionary advantage of jumbled contents of EVE clusters

We believe that the characteristic scrambling of EVE in PiRNA-like clusters when compared to their arrangements in the originating genome in both insects and shrimp is worthy of some contemplation. We propose that this phenomenon may have evolved because it prevents the easy re-establishment of complete, infectious viral genome sequences in the host genome by the process of recombination between EVE-cluster alleles. On the other hand, the ability of the EVE to produce RNA transcripts and give rise to vcDNA [11] would seem to open the possibility that rare recombination events might occur between infecting viruses and EVE products (RNA or DNA) and be an additional potential source viral variation. As far as we know, this possibility has not been explored in shrimp or insects. Is it possible, for example, that the origin of viruses in the “hybrid” family *Bidnaviridae* [22–24] might have originated by a fortuitous recombination event between distinct EVE cluster/ alleles that brought together EVE fragments from 2 different virus families into a novel, functional combination that led to the emergence of flacherie disease in the silkworm?

## CONCLUSIONS

Revelation of piRNA-gene-like clusters of EVE from specific viral types is worth further investigation. This may exemplify an advantageous evolutionary development that arose because collections of EVE from specific viral types into single linkage clusters would assure their transmission to offspring as potential “protective antiviral EVE packages” (PEVEP). It would also an evolutionary advantage to have these PEVEP located on different chromosomes to assure that maximum variation in PEVEP combinations would occur simply by random assortment of chromosomes during the production of gametes. If such PEVEP operated like alleles, additional variation would be possible via crossover during meiosis. Without viral genome fragmentation and scrambling (including reading direction) during the formation of these piRNA gene-like clusters, it might be possible that crossover between two suitable APP would occasionally re-establish a full, infectious viral genome or even give rise to new virulent types. Fracture and jumbling may have evolved to circumvent this possibility. However, it does not eliminate the possibly of pop-up, false-positive PCR amplicons that might arise from crossover between EVE that re-establish the sequence of a PCR target for a standard viral detection method. Nor does it eliminate the possibility of RNA and vcDNA arising from EVE might on very rare occasions be incorporated into novel functional entities (e. g., *Bidnaviridae*) or contribute to the evolution of infectious viruses via recombination events.

## MATERIALS AND METHODS

### Bioinformatics analysis of a draft *P. monodon* genome for IHHNV

The sequences of non-infectious IHHNV (GenBank record DQ228358) and infectious IHHNV (GenBank record AF218266) were used as the subject reference sequences for BlastN analysis (https://blast.ncbi.nlm.nih.gov/Blast.cgi) of a recently released whole draft genome (WDG) of the giant tiger shrimp *P. monodon* (GenBank accession no. JABERT000000000) [13]. Prediction of conserved DNA repeat sequences was determined using the Dfam database of repetitive DNA families (https://www.dfam.org/home) [25].

## Abbreviations

IHHNV: Infectious hypodermal and haematopoietic necrosis virus
IHHNV-A: non-infectious IHHNV Type A
cvcDNA: circular viral copy DNA
RT-PCR: reverse-transcriptase polymerase chain reaction
EVE: endogenous viral element(s)
piRNA: PIWI-interacting RNA(s)
RNAi: RNA interference

## Acknowledgements

This research project is supported by Mahidol University (Fundamental Fund: Basic Research Fund: fiscal year 2022) (Grant no. BRF1-054/2565) and National Center for Genetic Engineering and Biotechnology (BIOTEC), National Science and Technology Development Agency (NSTDA).

## Supplementary information

**Supplementary figure S1.**
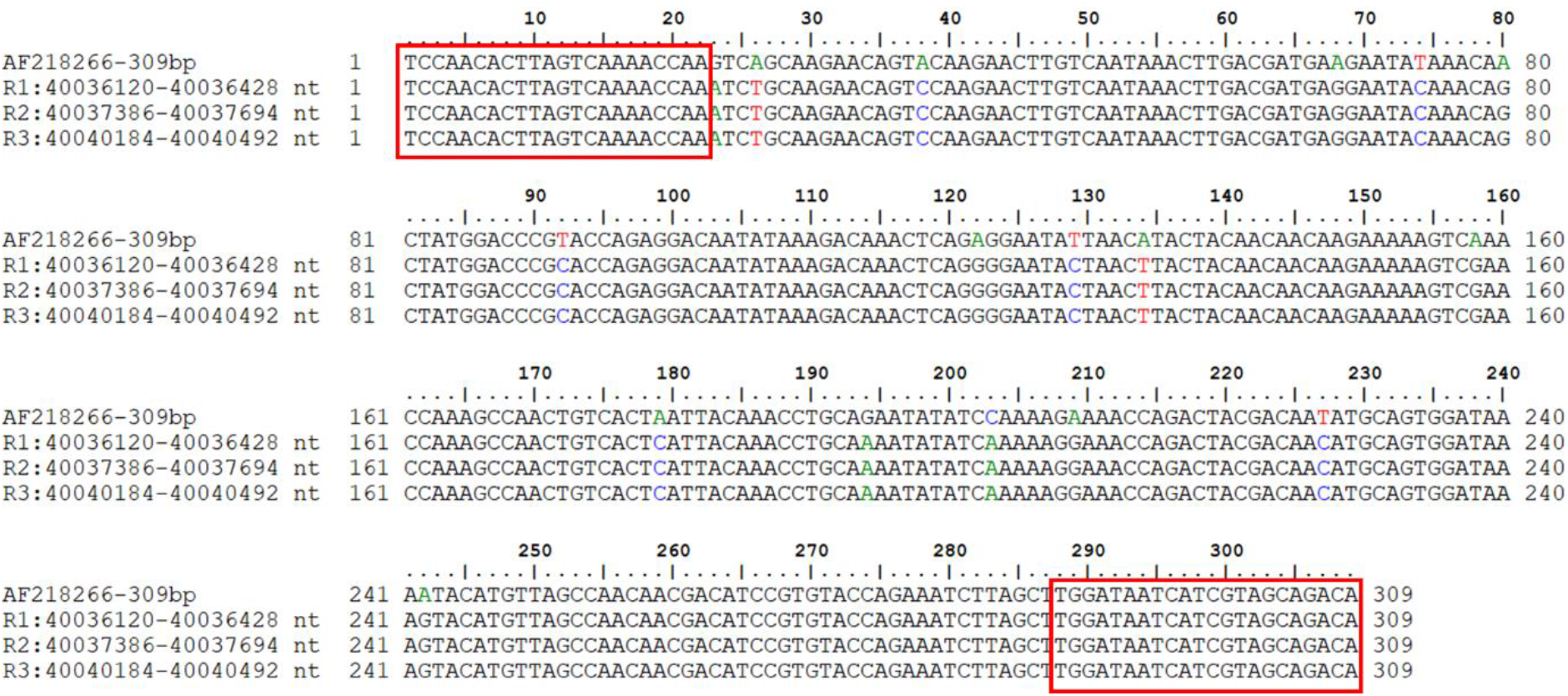
Multiple sequence alignment of potential 309 bp targets for the OIE- recommend IHHNV detection method from three repeats in IHHNV-EVE observed in LG 7 (R1-R3). The 309F/ R primer positions are indicated by red rectangles. The primer sequences and potential amplicon sizes match 100% to those expected for infectious IHHNV.

**Supplementary figure S2.**
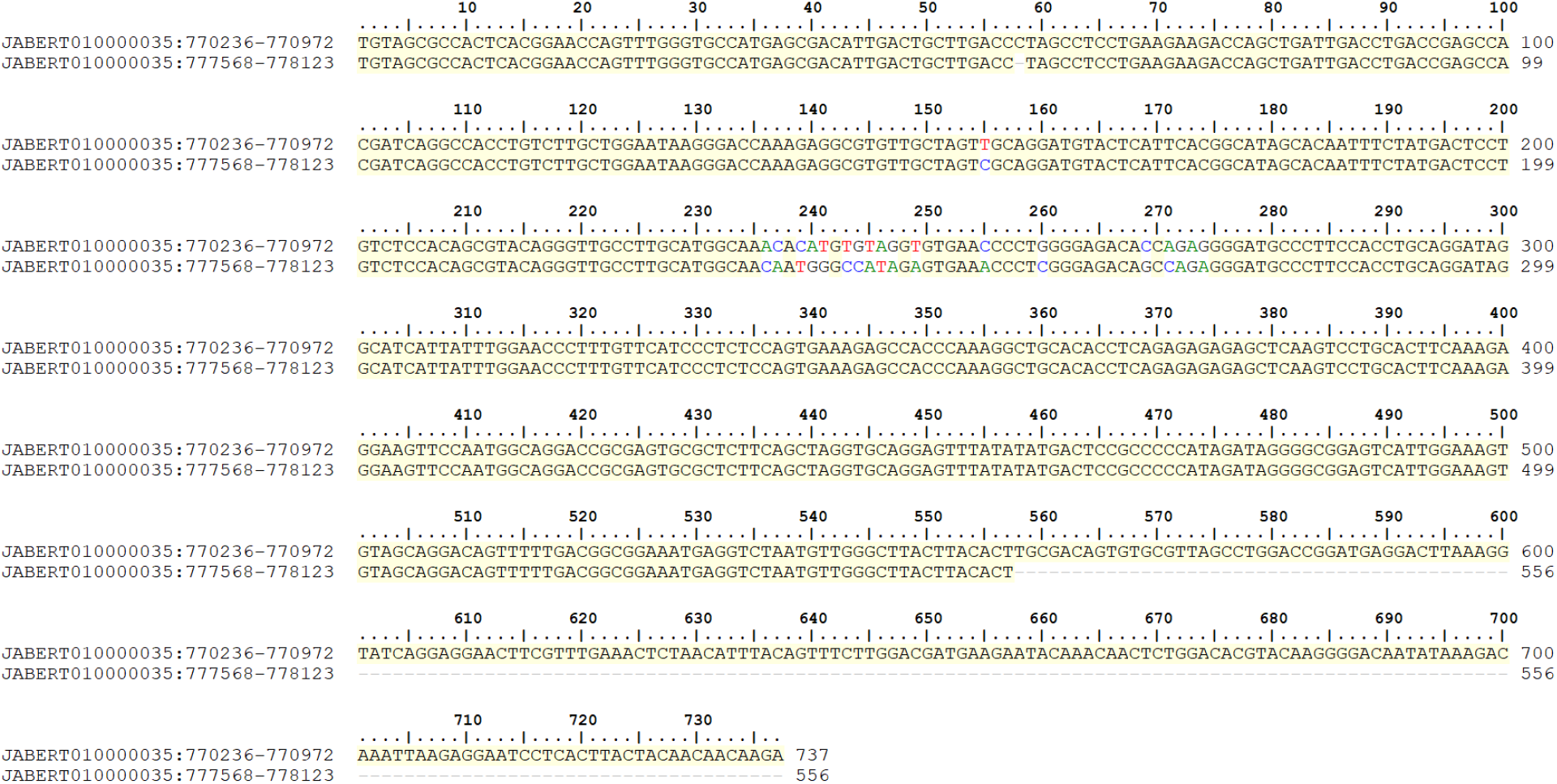
Alignment of the 561 bp repetitive sequences flanking the non-infectious IHHNV-EVE Cluster 1 located in LG35. The sequences are aligned in the same direction and share 98% identity (547/561 bp with 9/561 nucleotide gaps).

**Supplementary figure S3.**
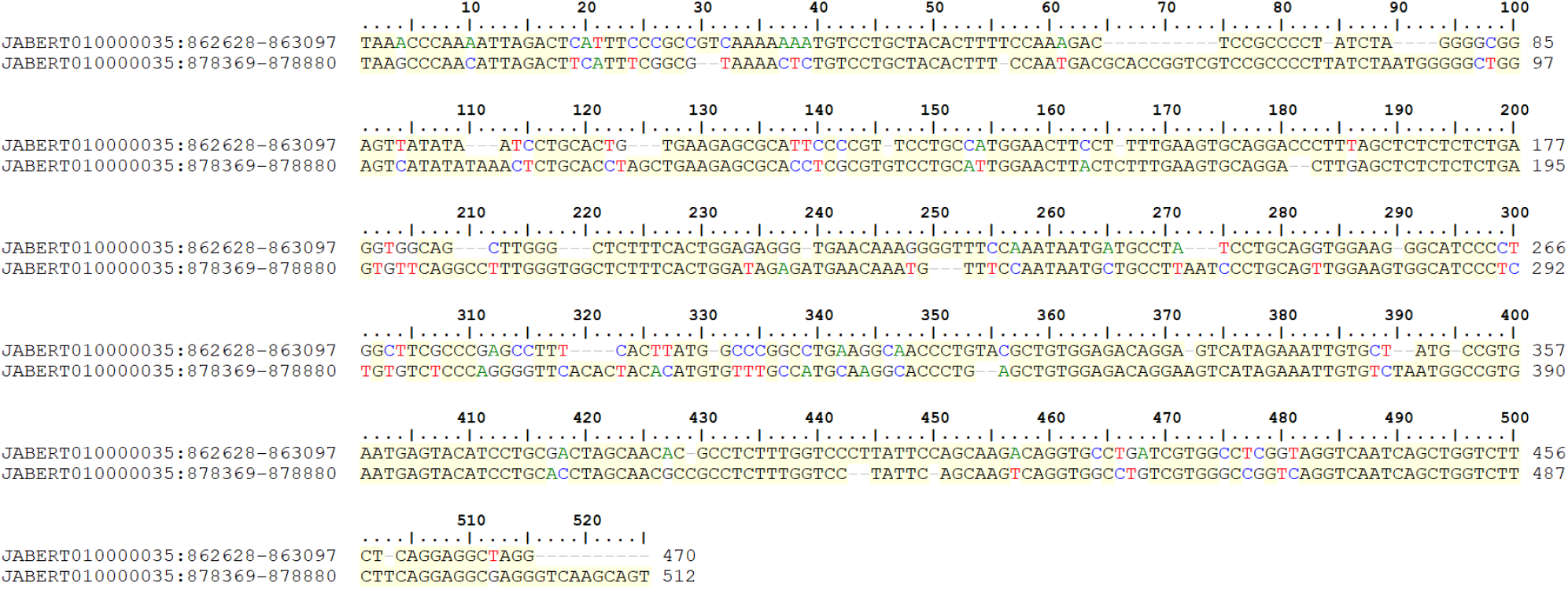
Alignment of the two 519 bp repetitive sequences flanking IHHNV-EVE Cluster 2 located on LG35. The sequences are aligned in the same direction and share 77% identity (400/519 bp with 66/519 nucleotide gaps).

**Supplementary figure S4.**
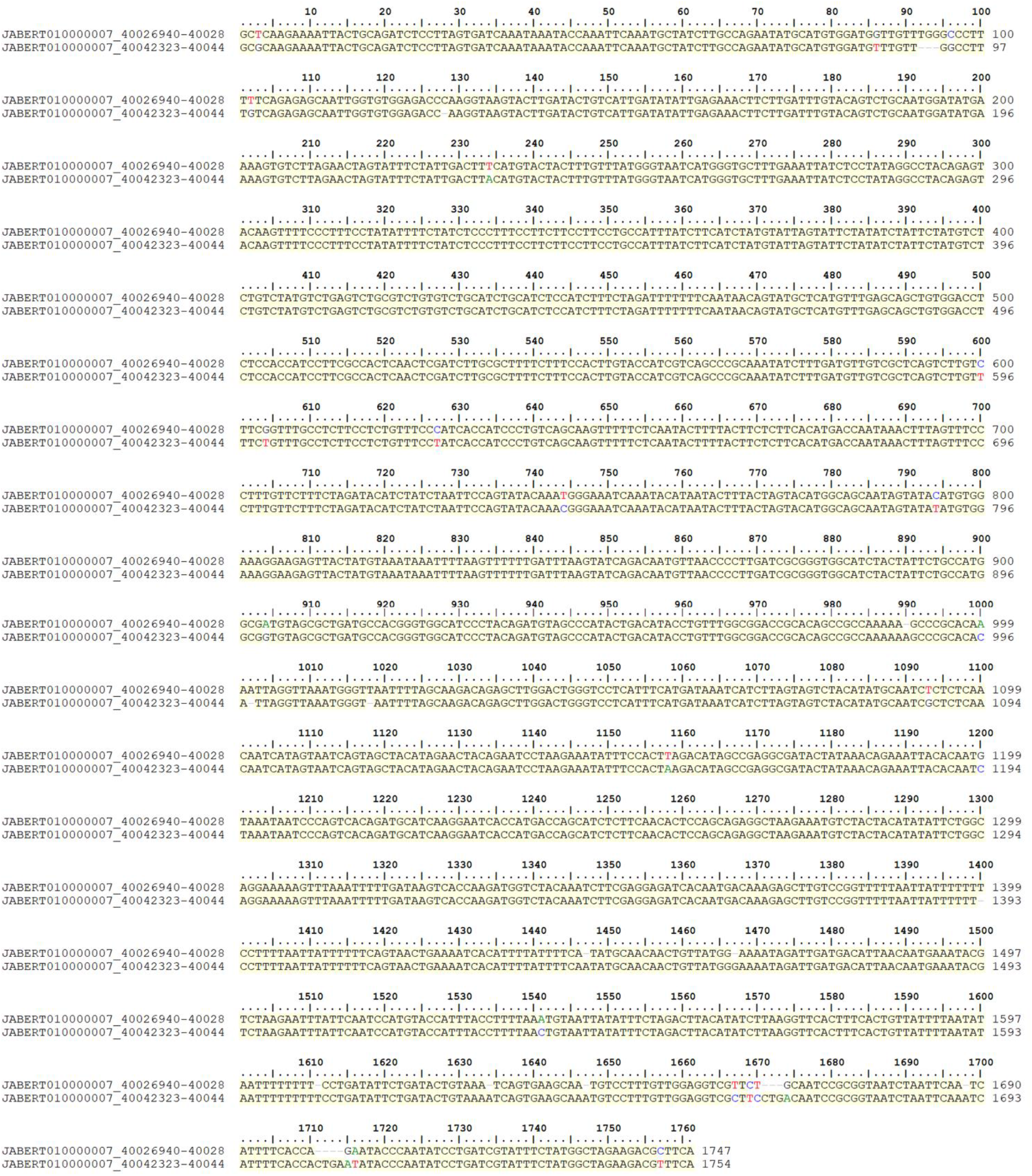
Alignment of the direct repetitive sequences flanking the infectious IHHNV-EVE Cluster 3 located on LG7. A total 1,758 bp were aligned and share 98% identity (1722/1758 bp with 21/1758 nucleotide gaps).

